# Cell-associated Transcriptional Alterations in the Retinal of Alzheimer’s Disease

**DOI:** 10.1101/2022.08.18.502974

**Authors:** Jennifer Ngolab, Adam Mark, Justin Buchanan, Shaina Korouri, Sebastian Priessl, Sara Brin Rosenthal, Allen Wang, Kathleen M. Fisch, Robert A. Rissman

**Author notes:** <u>Correspondence to:</u> Robert Rissman, Ph.D., Department of Neurosciences, UCSD School of Medicine, 9500 Gilman Drive, MTF 309, M/C 0624, La Jolla, CA 92093-0624.

## Abstract

Current approaches for studying pathologic changes in the retina associated with Alzheimer’s Disease (AD) remain heterogeneous, limiting the use of retinal amyloid-beta as a viable biomarker for AD. Transcriptomic profiling of the retina has provided cell-specific insight into AD progression in the brain yet is lacking in the retina. In this study, we implemented a non-biased approach through next generation sequencing to profile frozen archived retinal tissues from autopsy/pathologically confirmed AD and non-diagnosed cases (NonAD). A total of 37,211 nuclei were isolated from frozen retinal tissue punches originating from AD, and 31,326 were isolated from non-diagnosed cases. Gene expression patterns specific to the retinal region and major retinal cell types were represented in both tissue groups. AD-associated genes were differentially expressed in AD retinal glial cells, including microglia. A greater percentage of microglial nuclei from AD retinal nuclei expressed TYRO protein tyrosine kinase-binding protein (TYROBP) compared to nonAD retinal nuclei. However, compared to microglia from single retinal cell datasets from elderly non-diseased individuals, TYROBP expression is highly expressed in the single cell data set, indicating TYROBP transcripts reside within the cytoplasm. However, other AD-associated genes were differentially expressed in AD nuclei such as DOCK2, PICALM, and PLCG2 compared to non-diseased single-cell microglia, implicating a role of these genes in the AD retina. To summarize, we extracted a high number of nuclei from frozen retinal tissue that retain specific gene markers for cell classification and highlighted candidate AD-associated genes in retinal microglia that may be viable in future AD retinal studies.

## Introduction

Due to their shared embryonic origin and similar neuronal architecture, the photoreceptive retina is considered an outgrowth of the brain and therefore hypothesized to be susceptible to similar neurodegenerative processes ^1,2^. In the case of AD, retinal thinning and vascular alteration occurs in early and late-stage AD patients compared to age-matched controls ^3^. Clinical retinal imaging studies and post-mortem histochemical analysis of retinal tissue implies that amyloid-beta deposits accumulate in cases with high brain amyloid-beta ^4,5^. The potential of using the retina as a non-invasive biomarker could further aid in a faster and more comprehensive neurodegenerative diagnosis alongside other established hallmarks. An international committee has been formed to standardize protocols related to AD-associated retinal research, yet current data are limited in interpretation due to variation in hardware and laboratory techniques ^3^. Retinal alterations are indicative of retinal disease and normal aging ^6,7^. To address these confounding variables, other diagnostic parameters based on neurodegenerative biological pathways can further differentiate diseased from temporal-based retinal changes.

While retinal amyloid-beta deposition has been highly investigated in the AD retina ^8–14^, other hallmarks of neurodegeneration have been detected in the AD retina through histological analysis. Two separate studies identified tau accumulation within AD postmortem retinal tissues ^10,15^. Gamma-synuclein accumulated outside the ganglion cell layer in post-mortem retinas from AD cases, suggesting that alteration of synuclein occurs in AD ^16^. Retinal amyloid-beta deposition and the presence of phosphorylated tau coincided with increased astrogliosis and microglial reactivity, with increased TREM2 transcript identified through fluorescent in situ hybridization ^17^. These data suggest neurodegenerative states such as AD alter the glial cells regardless of the location of the neural tissue. Unlike what is observed in the brain, TREM2 expression failed to colocalize with microglia in the retina. The authors recognize that technical issues such as tissue preparation may impede microglial TREM2 detection. While histology provides region-specific assessment of tissue, additional experiments conducting cell-population analysis may capture a unique microglial transcriptional profile associated with AD diagnosis.

The rapid implementation of next-generation sequencing in ocular science has revealed much about the eye’s molecular profile, distilling the retina’s complex nature. Developing a comprehensive profile of the retinal sub-populations has identified cell-specific molecular changes in retinal diseases ^18^. The retina is highly susceptible to post-mortem degradation, as decreased MALAT1 expression in rod photoreceptors correlated with longer post-mortem times ^19^. Most studies isolated whole cells from retinal tissue extracted 6 hours post-mortem, which may be inaccessible to many researchers. Fortunately, nuclei from frozen retinal tissue yielded similar transcriptomic profiles to that of single-cell RNA-seq, indicating that transcriptional profiling can be implemented on archival tissues ^20^. Studies using archival brain tissues have identified disease-associated subpopulations of cells through transcriptomic profiling ^21–23^. The ability to analyze the nuclei of frozen archival tissue allows for comprehensive analysis between previously identified pathological and morphological hallmarks and potential disease-related transcriptional profiles. Extending next-generation analysis to the AD retina may provide additional biological insight into AD-mediated retinal changes.

While total transcript can be extracted from archival tissue using single-nuclei RNA-seq, detecting activation of state-associated genes is inconsistent between brain studies. On one hand, disease-activated genes were preserved in frozen microglial nuclei ^24,25^. However, a meta-analysis of publicly available microglial datasets indicates that microglial activation genes may be depleted in single-nuclei RNA-seq datasets, including genes involved in AD ^26^. However, as evident in immunohistochemical studies, disease-associated transcriptional changes in the brain may not be evident in the retina and the genes that indicate retinal microglia activation ^17^.

This study aimed to determine if AD-associated transcriptomic changes documented in the brain also occur in the retina. To investigate this, we have isolated nuclei from archived retinal tissue from AD cases and NonAD cases. A total of 37,211 nuclei were isolated from two frozen retinal tissues from histologically confirmed AD cases, and 31,326 nuclei were extracted from two non-diagnosed cases. Unsupervised clustering of the nuclei’s gene expression profiles generated clusters corresponding to the seven major retinal cell populations. Genes associated with specific retinal regions were preserved, with no significant shifts in expression between AD and NonAD nuclei. The expression of genes associated with AD varied within retinal glia cells, with significant differences in the expression found in APOE, HLA-DQA1, HLA-DRB1, TYROBP, and CLU in microglia and APP and APOE in Muller cells. We aligned our retinal single-nuclei data with a single cell database derived from aged undiagnosed retinal tissues to determine if flash freezing may have affected our data. However, when investigating known markers of microglia activation, genes identified previously between our AD and NonAD samples were highly expressed in the healthy dataset. However, other AD-associated genes, such as GAB2, PICALM, DOCK2, PTK2B and PLCG2 were differentially expressed, suggesting other AD genes may be upregulated in the retina. Together, these data may provide further insight into the molecular changes that underlie the morphological changes to the retina in AD.

## Results

### Characterization of retinal nuclei extracted from frozen tissue

Nuclei were harvested from 4 mm tissue punches extracted from the peripheral region and macular fovea of frozen retinal tissue originating from pathology confirmed AD cases (n = 2) as well as non-diagnosed late adult retinal samples (n=2), resulting in eight libraries. After filtering out low-quality nuclei, 37,211 nuclei were isolated from AD tissues and 31,326 from NonAD tissue. Clustering analysis through reciprocal PCA grouped all nuclei into 31 clusters (Fig. 1a). Nuclei from every sample were represented in each cluster, with most clusters exhibiting a bias for nuclei from AD samples (Fig. 1b). A curated list of known retinal cell gene markers (PDE6A for rod photoreceptors, ARR3 for cone photoreceptors, NETO1 for Bipolar cells, ONECUT2 for horizontal cells, RGR for Muller glia, ELAVL2 for retinal ganglion cells (RGC), ATP1A2 for astrocytes, GAD1 for amacrine, and HLA-DRA for microglia) was used to assign clusters to a preliminary retinal cell type (Fig. 1b, Supplemental Table 1). Cluster 22 did not express any of the aforementioned markers but upon further review, was clustered into the amacrine cell identity, due to the proximity of the cluster to other amacrine-identified clusters (Fig. 1a). Conversely, several clusters expressed more than one gene marker, such as cluster 26 containing a sizable percent of nuclei expressing PDE6A and RGR (Fig. 1b). All representative gene markers were expressed in the AD, and Non-AD samples (Fig. 1c). Differential gene expression analysis revealed additional cell-specific markers specifically expressed in the putative cell clusters, which are highly involved in pathways specific to the retinal cell type (Fig. 1d). We quantified the proportion of each cell type within the diseased and non-diseased tissues. While both samples had a comparable number of cone photoreceptors (NonAD: 1891, AD: 1916), 10,612 rod photoreceptor nuclei were quantified in the NonAD condition (Fig. 1f, Table 1) while a greater number of astrocytes, microglia and horizontal cells were grouped in the AD group (Fig. 1g, Table 1).

**Fig. 1.**
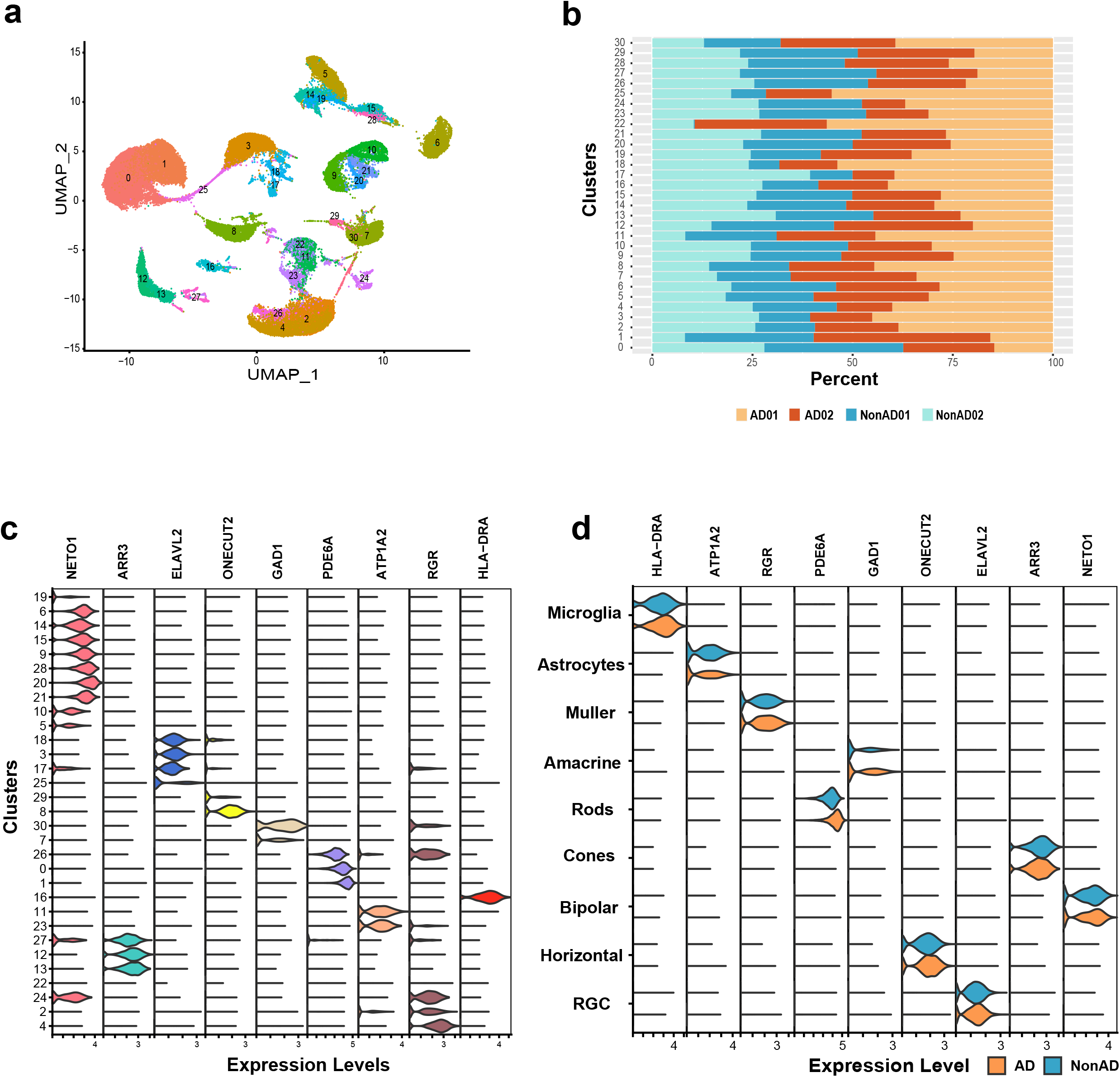
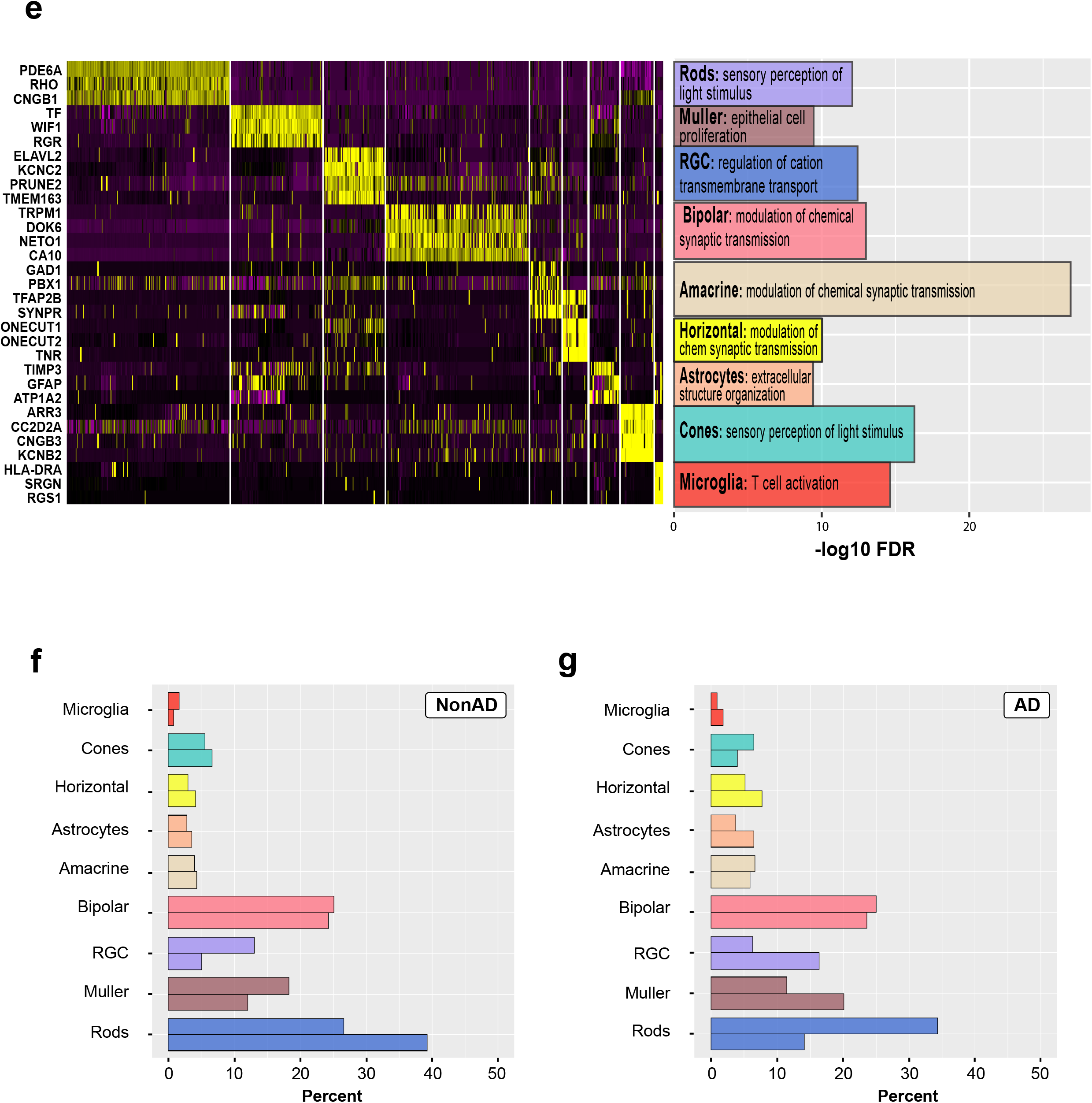
Characterization of frozen retinal samples through unsupervised clustering. **a)** Unsupervised clustering identifies 31 distinct clusters. **b)** Proportion of samples composing each cell cluster. **c)** Assigning cell identity through gene markers. **d)** Expression of gene markers between disease status. Orange: AD, Teal: NonAD. **e)** Heatmap highlighting other retinal marker genes in putative cell-type clusters. The bar graph indicates the highest pathway retrieved from gene enrichment analysis. The - log10 of the False Discovery Rate of the highest enriched term is plotted. Segregation of **f)** NonAD and **g)** AD nuclei into cell type. Data is mapped as a percentage of total nuclei. RGC: Retinal Ganglion Cells, UMAP: Uniform Manifold Approximation and Projection

**Table 1.**
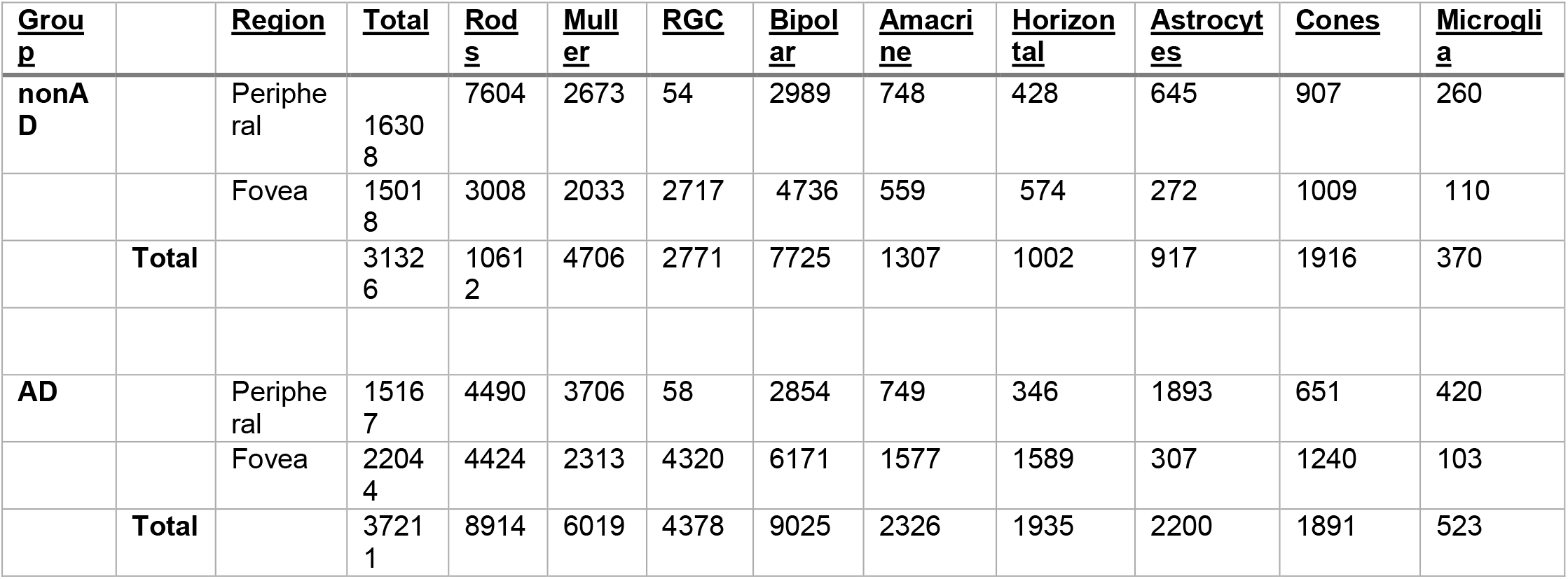
Number of nuclei extracted, tabulated by source, region and cell type. RGC: Retinal ganglion cells. AD: Alzheimer’s Disease retina, nonAD: non-diagnosed retina

| <u>Group</u> |  | <u>Region</u> | <u>Total</u> | <u>Rods</u> | <u>Müller</u> | <u>RGC</u> | <u>Bipolar</u> | <u>Amacrine</u> | <u>Horizontal</u> | <u>Astrocytes</u> | <u>Cones</u> | <u>Microglia</u> |
| --- | --- | --- | --- | --- | --- | --- | --- | --- | --- | --- | --- | --- |
| **nonAD** |  | Peripheral | 16308 | 7604 | 2673 | 54 | 2989 | 748 | 428 | 645 | 907 | 260 |
|  |  | Fovea | 15018 | 3008 | 2033 | 2717 | 4736 | 559 | 574 | 272 | 1009 | 110 |
|  | **Total** |  | 31326 | 10612 | 4706 | 2771 | 7725 | 1307 | 1002 | 917 | 1916 | 370 |
| **AD** |  | Peripheral | 15167 | 4490 | 3706 | 58 | 2854 | 749 | 346 | 1893 | 651 | 420 |
|  |  | Fovea | 22044 | 4424 | 2313 | 4320 | 6171 | 1577 | 1589 | 307 | 1240 | 103 |
|  | **Total** |  | 37211 | 8914 | 6019 | 4378 | 9025 | 2326 | 1935 | 2200 | 1891 | 523 |

We next sought to confirm region-specific differences within our samples. Differential gene analysis localized region-specific genes such as SNCG, NEFL, and PVALB to fovea samples. In contrast, periphery-associated genes such as SAG and SLC1A3 were enriched in samples isolated from the periphery (Supplemental Fig. 1a). There were differences in the percent of nuclei expressing region-specific genes in AD and NonAD, such as the peripheral gene ID3 expressed in 43.7% of AD periphery nuclei as compared to 25.6% in NonAD nuclei (Supplemental Fig. 1b, Supplemental Table 1). However, the differences in region-specific genes between AD and NonAD, such as ID3 only trend towards significance (avg fold change = 1.25, p =0.011857). Taken together, these data indicate that the retinal gene expression profiles isolated from our frozen retinal samples replicate existing studies, suggesting that our dataset represents the molecular environment of the retina.

### Nuclei extracted from AD retinal tissue exhibit gene expression profiles

Because the sex of each case was correlated with disease status, the highest differentially expressed genes were sex-associated (Supplemental Fig. 2a). Other genes highlighted through differential gene expression include HSP1A, HSPH1, HILPDA, CABP5 and MMP2 (Supplemental Fig. 2b). Apart from HSPH1, these genes are highly expressed in NonAD nuclei, primarily in the photoreceptors. Of interest, CABP5 is associated with light sensitivity in rod photoreceptors. When factoring cell type identity, significant expression of HSPA1A, HSPH1, and HILPDA were identified in NonAD cells, namely the Amacrine and Bipolar cells, along with the photoreceptors (Supplemental Table 2). Interestingly, about 45% of AD Astrocytes express XIST, a non-coding RNA that is involved in the inactivation of the extra X chromosome and is highly expressed in female tissue (Fig. 2b, Supplemental Table 2).

**Fig. 2.**
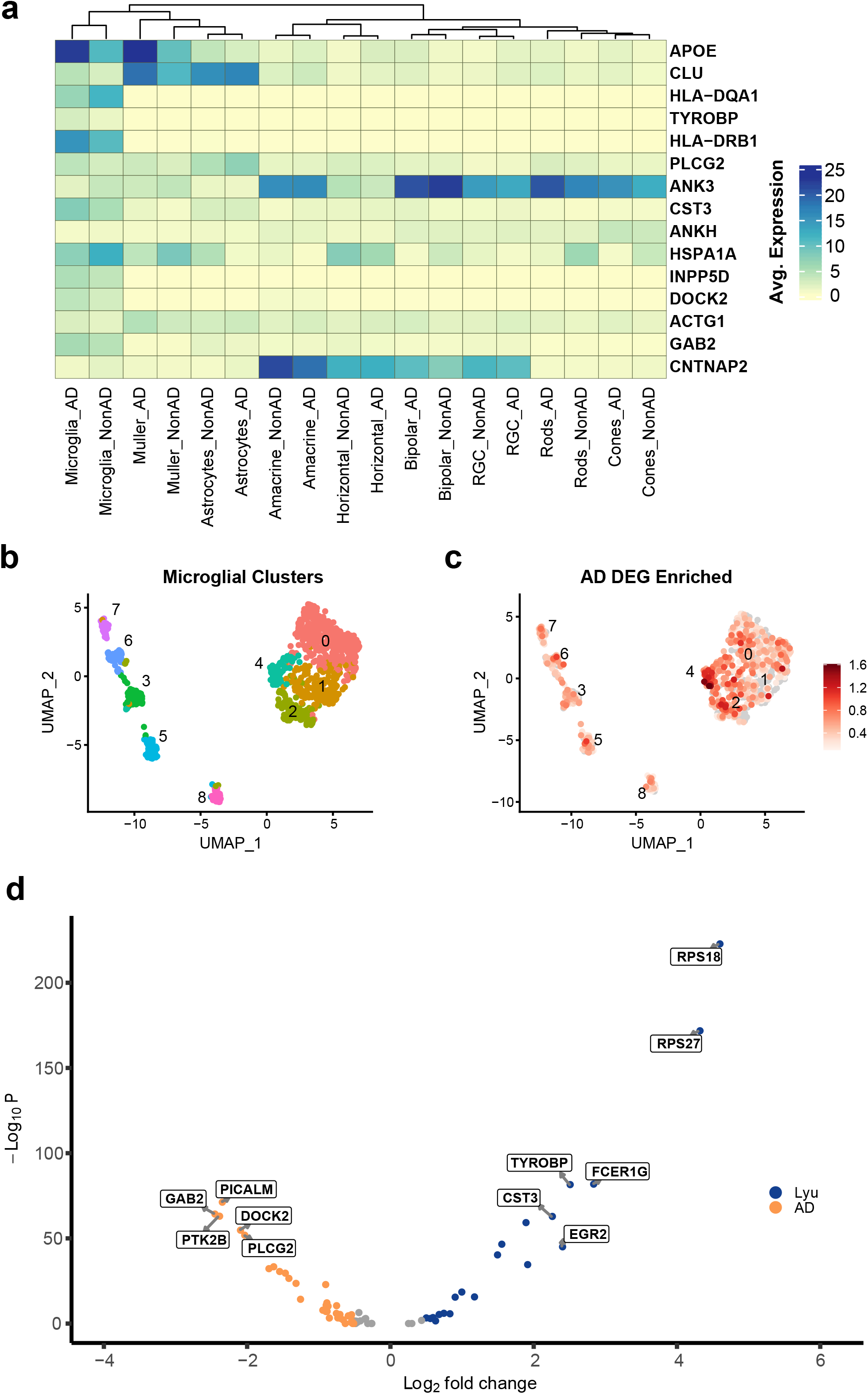
AD-associated genes are enriched in microglial nuclei. **a)** Average expression of AD-associated genes amongst cell type and disease state. **b)** Subpopulations of nuclei **c)** Abundance of AD-associated genes within each subpopulation of microglial nuclei. **d)** Volcano plot of differentially expressed genes between AD microglial nuclei and non-diseased age-matched microglial cells (Lyu). Log2 Fold changes are plotted against the log of the adjusted p-value. Colored dots indicate genes expressing a log fold change >=1 and an adjusted p-value of 1.0×10^-10. Orange dots originate from the AD microglia nuclei, and blue dots originate from the Lyu dataset. Labeled dots represent the highest expressed genes in each dataset.

### AD-associated genes localize to retinal microglia

To determine if increased brain amyloid-beta influences the neurodegenerative pathways in the retina, we examined the average gene expression levels of AD-associated genes derived from GWAS studies ^26,27^. Most AD-associated genes were contained within the glial cells of the retina, with the microglial nuclei exhibiting elevated levels of 8 out of the 13 AD genes differentially expressed in AD microglia compared to NonAD microglia (Fig. 2a). APOE was significantly expressed in AD Muller glia and microglia, with a slight but significant decrease observed in astrocytes (Muller: fold change = 2.85, p =0; Microglia: fold change = 2.22, p = 3.5 x 10-171, Fig. 2a, Supplemental Table 2). HSPA1A was highly expressed in most NonAD cell types except for RGCs (Fig. 2A, Supplemental Table 2). Microglial nuclei were extracted from the main dataset, and 893 nuclei were clustered to identify subpopulations of cells with a high concentration of AD-associated genes (Fig. 2b). Clusters 2 and 4 contained a high percentage of nuclei enriched in AD-Associated genes (33.3 and 78% of total cell in cluster) (Fig. 2c, Supplemental Table 2), with TYROBP differentially expressed in cluster 4 (fold change: 1.53, p = 6.63×10-11, Supplemental Table 2). Cluster 4 contains 50% of AD nuclei highly enriched with AD-associated genes, while only 28% of NonAD nuclei contain similar levels (Fig. 2d and Fig. 2e, Supplemental Table 2). While interesting and significant, the fold change between AD and nonAD cell types is less than 2 (Supplemental Table 2). Complications from our samples, such as age, sex, and sample size may also hamper our analysis.

### Comparison of single-nuclei RNA analysis with previously validated single-cell RNA datasets

To further support our observations and verify the validity of our data, we compared our single retinal nuclei AD data to a publicly available retinal single-cell dataset extracted from ophthalmologic confirmed healthy retinas from elderly male individuals, which will be referred to by ‘Lyu’ ^27^. The nuclei of both datasets were clustered by cell type, and each dataset expressed a similar amount of each marker (Supplemental Fig. 3c, Supplemental Table 2). Furthermore, genes enriched in the Lyu cell types were similarly expressed in the corresponding AD dataset (Supplemental Fig. 3, Supplemental Table 2). We subset the microglial cells from both AD and the Lyu dataset, resulting in a total of 832 cells. Differential gene expression analysis revealed that TYROBP was highly expressed in the Lyu dataset, along with FCER1G, CST3, and EGR2 (Fold change >2, Fig. 2d). Conversely, AD-associated genes differentially expressed in the AD dataset included GAB2, PICALM, DOCK2, PTK2B, and PLCG2 (Fold change >2, Fig. 2d). While expression was higher in the Lyu data set (fold change = 0.90, padj = 3.20×10-16), the percentage of cells expressing APOE was similar between the two datasets (Lyu: 89.5%, AD: 92.9%, Supplemental Table 2). While genes associated with microglial activation were lowly expressed in our nuclear dataset, other AD-associated genes were differentially expressed in microglia, suggesting that other genes involved in AD are implicated in pathways leading to AD-mediated retinal degeneration (Supplemental Fig. 3d).

## Discussion

Our study aimed to identify molecular changes that may influence retinal restructuring observed in Alzheimer’s Disease patients. We isolated 37,211 nuclei from two frozen AD retinal tissues and 31,326 nuclei from two frozen non-diagnosed cases, indicating that freezing does not impair the number of intact nuclei. All seven major retinal cell populations were identified using known gene markers in our sample, including microglia ^28^. The localization of known AD-associated genes to the retinal microglia cells further support the involvement of microglia in AD-mediated retinal pathology. To correct for the variations in our single-nuclei dataset, we compared our AD retinal dataset to a single-cell retinal dataset derived from age and sex-matched non-diagnosed cases ^27^. While TYROBP and other AD-associated genes lowly expressed in our NonAD sample were enriched in the non-diseased Lyu single cell dataset, other AD-associated genes were differentially expressed in the AD microglia of our retinal nuclear dataset. We demonstrate the utility of nuclei extraction from archived retinal tissue from cognitively compromised individuals, and potentially identifying transcriptional profiles unique to retinal AD.

Our primary focus was to confirm all retinal cell type gene expression profiles from frozen retinal tissues from pathologically diseased cases. Multiple transcriptomic studies employed a trained ophthalmologist to visually assess the retina before tissue harvesting to account for any retinal diseases that may influence the data ^20,27,29^. Unlike previous studies, our tissues originated from pathology-confirmed neurodegenerative cases originating from the UCSD ADRC and lack any ophthalmologic examination. A high number of nuclei were extracted from frozen AD and nonAD retinal tissue, with a sizable number of genes read (Table 1, Supplementary Table 1). Similar retinal nuclei studies isolated far fewer nuclei despite using a larger diameter punch but a high number of reads per nuclei due to the technology used to capture and sequence the nuclei ^20^. Interestingly, our numbers were consistent with other single-cell retinal RNA-seq studies using larger punch diameter. Therefore we conclude that our methodology of isolating from diseased retinal tissue does not compromise the quality of the isolate. Likewise, our non-diseased sample originates from cases with no confirmed brain pathology. Future studies seeking to conduct brain-retina comparison studies should plan on obtaining ophthalmologic, cognitive clinical, and pathological data to better assess the interactions between the two organs. We acknowledge the technical issues of the current study, and therefore our conclusions with the current data may be limited. Because of the high yield of nuclei extracted from the study, we anticipate building upon these findings by analyzing samples from our AD retinal tissue archive.

While we aimed to keep all demographic variables between cases as similar as possible, it was difficult to obtain non-diseased tissue that shared characteristics with our ADRC samples. The sex of each case limits our gene expression analysis between disease states. We observed bias towards sex-differential genes (Supplemental Fig. 1a). The addition of the opposite sex to each disease group may reduce the influence of sex-specific genes in our differential gene expression analysis. Curiously, the sex-specific gene XIST was expressed in peripheral AD astrocytes. The AD astrocyte nuclei that express XIST also express UTY, indicating that no contamination has occurred (Supplemental Fig. 2b). However, it is possible that this could be due to an unwanted technical error during processing or alignment. Nuclei extraction through homogenization can result in ambient RNA, which can result in RNA contamination and obscure downstream analysis ^30^.

Despite the variations amongst our nuclei dataset, AD-associate genes related to glial cells were differentially expressed in AD samples. HSPA1A is an immediate early gene involved in responding to stress and injury that are implicated in the slow progression of neurodegenerative disease ^31^. Interestingly, a greater percent of NonAD nuclei express HSPA1A, especially in the retina neuronall cells, suggesting that the expression is downregulated in AD (Fig. 2a, Supplemental Table 2). An increase of TYROBP expression was also observed in AD microglia compared to NonAD microglia (Fig. 2a). The increase of TYROBP AD microglia indicates a transition from a homeostatic glial cell to a disease-activated microglia ^32,33^, suggesting that AD-associated activation of microglia may occur after high levels of brain amyloid. Previous histochemical analyses have also identified the disease-activated microglial phenotype in AD retina, albeit with TREM2 ^17^. While TYROBP is known to interact with TREM2, a recent study in mice suggests an interaction between TYROBP and APOE in microglia results in reduced amyloid and an increase of phosphorylated tau ^34^. The presence of phosphorylated tau has been identified in AD postmortem retinal tissue, specifically in the retinal periphery where our AD microglia were isolated ^10,15^. Immunohistochemical analysis detecting phosphorylated tau and/or DAP10 expression in AD retinal tissue would further investigate this possible relationship.

To correct for the demographic discrepancies in our study, we supplemented our AD nuclei dataset with a public retinal single cell dataset extracted from elderly male individuals with no obvious abnormalities on the retina ^27^. Single nuclei RNA-seq datasets have corroborated their whole cell counterparts, with studies reporting an overlap of genes expressed ^35^. Despite reports indicating compatibility between single nuclei and single-cell datasets, expression profiles from the single cell RNA-seq contain additional transcripts in the cytoplasm, such as mitochondrial genes, which may offset differential gene analysis ^36^. Detection of genes associated with disease has been less successful through snRNA-seq, suggesting that key disease markers localize to the cytoplasm. Key genes associated with microglia activation, such as APOE, CD74, and CST3 were depleted in single nuclei brain microglia samples, leading the authors to conclude that single nuclei RNA-seq should not be used for detecting disease-related subpopulations ^26^. Indeed, we observed that CST3 was differentially expressed in the whole cell database (Fig. 2d, Supplemental data 3). However, even though APOE was proportionally higher in the single cell dataset, it is expressed in a large percentage of microglial cells (Supplemental Table 2). Furthermore, CD74 was not found to be differentially expressed yet present in both datasets, suggesting that certain disease-associated genes may be preserved in archived retinal tissues (Supplemental Table 2). It is important to acknowledge that the single cell data set used in this study represents a fraction of retinal transcription profiles. Organ-specific cell reference atlases are available to compare new datasets to publicly available datasets. A reference atlas composed of 33 publicly available retinal single-cell RNA-seq datasets was recently made available ^37^. The ability to compare new retinal data to an atlas of healthy retinal cells can further identify subpopulations relevant to the diseased retina.

Genes differentially expressed in our AD nuclei dataset include GAB2, PICALM, DOCK2, PTK2B, and PLCG2 (Fig. 2d). Most highlighted genes are essential for microglial cytokine release and endocytosis, as well as implicated in amyloid beta pathogenesis ^38,39^. Single nucleotide polymorphisms on the GAB2 gene are linked with a higher risk of late-onset Alzheimer’s Disease in APOE ε4 carriers ^40^. Both AD samples in our study confer an ε4 genotype. The effects of these highlighted genes are documented in the brain yet are unknown in the retina. Mice retinal transcriptomics failed to detect the mouse equivalent of DOCK2 ^41^. Further analysis of human retinas to confirm the presence of these genes would further support our initial findings.

This study assessed the feasibility of detecting retinal-specific AD-associated transcriptional changes in post-mortem autopsy-verified AD retinal tissues. Investigating the role of genes such as GAB2, PICALM, DOCK2, PTK2B, and PLCG2 in the AD retina could potentially feed into the development of retinal diagnostic markers and inform of the relationship between brain and retina AD pathology. Further assessment of how gene expression in retinal glial gene changes over AD progression may offer insight into distinguishing retinal changes induced by age from disease-instigated retinal alterations.

## Materials and Methods

### Human Samples

Both ocular globes were obtained about 6-12 hours after death and dissected immediately upon retrieval. Post-mortem AD retinal tissues were obtained from longitudinal AD cohort autopsies at the Shiley Marcos Alzheimer’s Disease Research Center Neuropathology Core at the University of California, San Diego. As a technical control, retinal tissues from clinically undiagnosed cases were obtained from the San Diego Eye Bank. Demographic information is listed in Table 1. The anterior segment of the right ocular globe was removed, and the resulting posterior pole was submerged in 4% PFA and stored at 4 C until use. The retinal tissue from the left ocular globe was dissected and portioned into inferior, nasal, superior, and temporal quadrants using parameters and previously validated protocol ^15^. Each quadrant was subjected to snap freezing and stored in −80 C until use.

### Nuclei Harvesting and Sequencing

A 4mm punch was extracted at the superior peripheral region and the fovea from snap-frozen retinal tissue using a disposable skin biopsy punch tool. Punches were kept at −80 C until isolation. Nuclei were extracted from each sample using a dounce homogenizer under chilled conditions. The dounce homogenizer was primed with NIM-DP (.25M Sucrose (S1888, Sigma), 25 mM KCl (AM9640G, Invitrogen), 5 mM MgCl2 (194698, Mp Biomedicals Inc), 10 mM Tris-HCl, pH 7.5 (15567027, Thermos Fischer Scientific), Molecular biology water (46000-CM, Corning), 25 mM DTT (D9779, Sigma), 1X Protease inhibitor (Sigma, NC0969110), 0.2 U/ μl RNasin (Promega, PAN2111)) buffer before each tissue punch was processed. Each tissue punch was resuspended in NIM-DP-L (1X NIM-DP, 0.1% Triton X-100 (Sigma, T8787-100ML)) buffer before being transferred to the dounce homogenizer. Samples were homogenized with a loose plunger until the tissue had been broken into small pieces, then homogenized with a tight plunger until the solution was uniform with no obvious particles. A 30 μm CellTrics filter (Sysmex) was used to filter the resulting homogenate before. Nuclei were spun down using a swinging bucket centrifuge (1000 x rcf for 10 min at 4°C). The pellet was resuspended in 600 μl of Sort Buffer (1 mM EDTA (Invitrogen, 15575020), 0.2U/ul RNasin (Promega, PAN2111), 2% BSA in PBS (Sigma, SRE0036-250ML; Corning, 21-040-CV)), then supplemented with DRAQ7 (Cell Signaling Technology, #7406S). 60,000 nuclei were sorted using the SH800S (Sony), into a Lobind tube (Eppendorf) with 50 μl of Collection Buffer (1 U/ul RNasin (Promega, PAN2111), 5% BSA in PBS (Sigma, SRE0036-250ML; Corning, 21-040-CV)). Nuclei were pelleted in a swinging bucket centrifuge (1000 x rcf for 15 min at 4°C) and resuspended in reaction buffer (RNase inhibitor (0.2 U/μl) and 1% BSA (Sigma-Aldrich, SRE0036) in PBS). An aliquot of the suspended solution was mixed with Trypan blue and loaded into a hemocytometer to assess nuclei concentration. Single-nucleus GEM reactions and libraries were generated using the Chromium Next GEM Single Cell 3’ Reagent Kits v3.1 kit (10x Genomics) as per manufacturer specifications (v3.1_User_Guide) with the following parameters: cDNA libraries were generated using 12 PCR cycles for downstream library processing, and 10 PCR cycles were used to generate sufficient library quantity of good complexity for sequencing. The final library concentration was assessed using the Qubit dsDNA HS Assay Kit (Thermo Fisher Scientific), and fragment size was checked using TapeStation High Sensitivity D1000 (Agilent). Libraries were sequenced using a NextSeq 500 or NovaSeq 6000 (Illumina) using these read lengths: read 1, 28 cycles; index read 1, 10 cycles; index read 2, 10 cycles; read 2, 90 cycles.

### Data Analysis

Sequence reads were aligned and sorted into feature-barcode matrices through the 10X CellRanger pipeline. Analysis of nuclei was conducted using the Seurat package in R ^19^. Integration of the snRNA-seq datasets was conducted through reciprocal PCA. Visualization of nuclei was done using UMAP ^20^. Differential gene expression to identify the retinal region, status, and cell-specific gene markers was conducted using the general linearized model MAST ^21^. Gene enrichment analysis was conducted using the clusterProfiler package ^22^. Integration of publicly available datasets with the current studies dataset was merged using the LIGER algorithm. Heatmaps were generated using the *ComplexHeatMap* package ^23^, volcano plots were generated using the *EnhancedVolcanoPlot* package ^24^, and additional visualizations were conducted using the scCustomize package ^25^.

## Supporting information

Description of Supplemental Files

## Acknowledgements

We thank Jeffrey Metcalf and other members of the Rissman lab for technical support for this project. We also thank K. Jepsen and the UCSD IGM Genomics Center for sequencing the snRNA-seq libraries. This work is funded by NIH/NIA grants AG070595, AG0518440, AG051848, AG058533 and funding to the Shiley Marcos ADRC Neuropathology core from P30 AG062429 (to RAR). This work was also funded by the Altman Clinical and Translational Research Institute (ACTRI) grant # UL1TR001442. Work at the UCSD Center for Epigenomics was supported by the UC San Diego School of Medicine.

## Data Availablility

Single-nuclei data from AD and nonAD retinal tissues will be available upon request. Single-cell retinal data used in this study can be accessed through GEO accession number GSE155288.

---

**Supplementary Fig 1.**
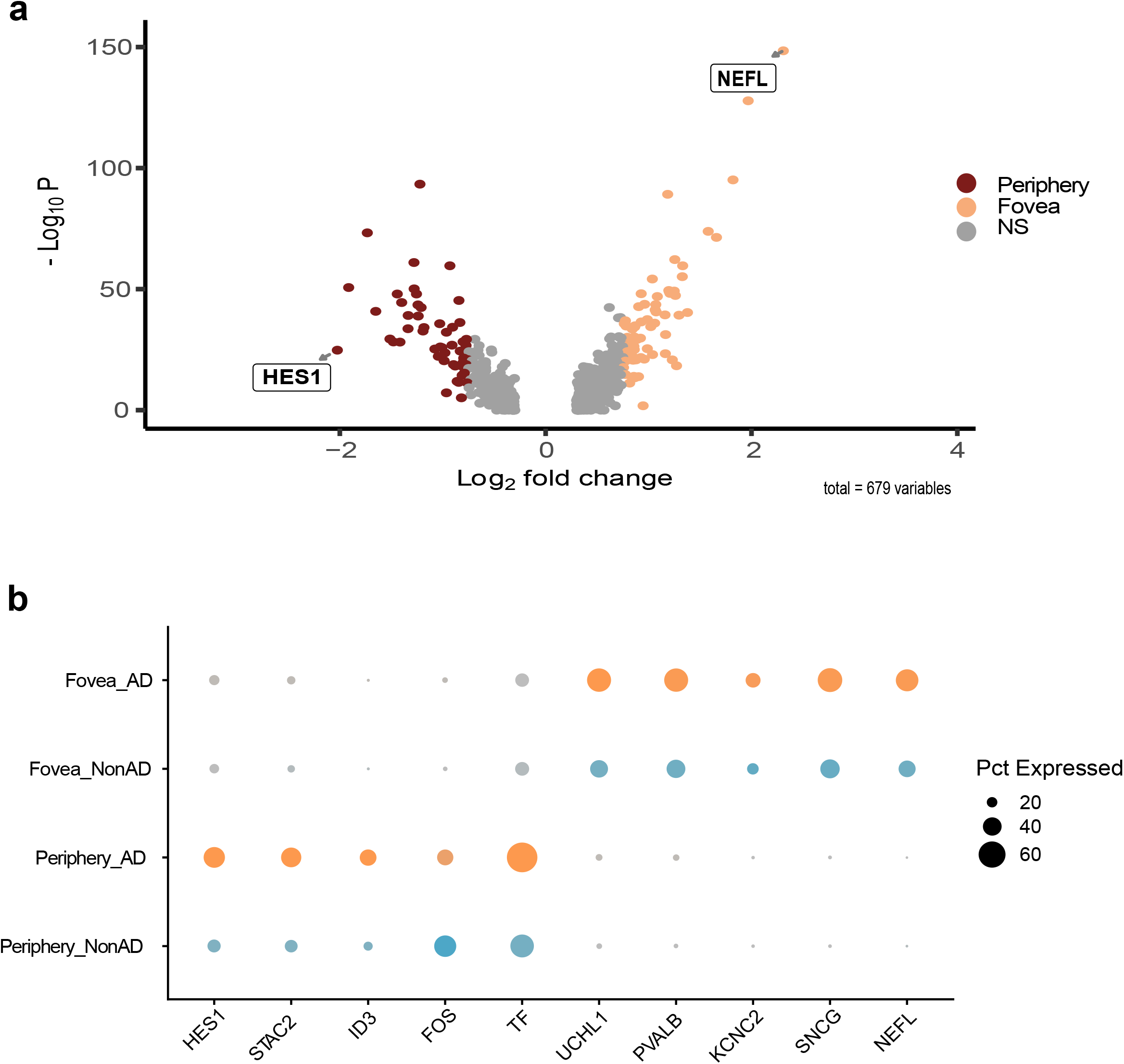
Transcriptional characterization of retinal regions between AD and NonAD single retinal nuclei. **a)** Volcano plot of differentially expressed genes between fovea and periphery. The y-axis indicates significance while the x-axis shows the magnitude of expression. Grey dots indicate non-significant genes. Blue dots indicate genes deemed significant yet are lowly expressed. Green dots indicate genes that meet or surpass the log-fold threshold (0.5). Red are genes that are expressed highly and are considered significant. A total of 1759 variables were considered in creating the graph. **b)** Expression of regional-specific genes between AD and NonAD samples. Blue = AD samples, Grey = NonAD samples.

**Supplementary Figure 2.**
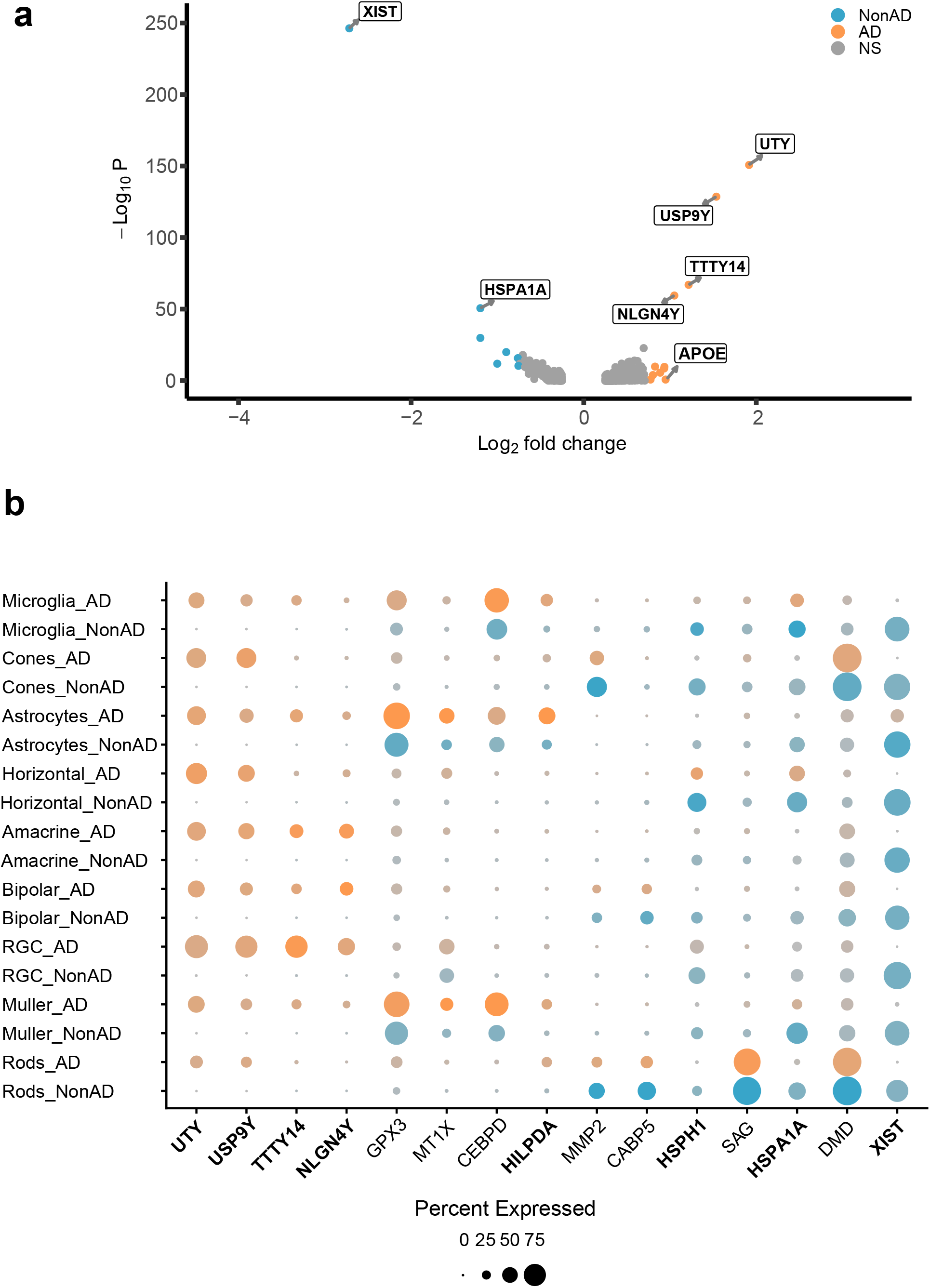
Slight dysregulation of gene expression within AD samples. **a)** Volcano plot of differentially expressed genes between NonAD and AD samples. The y-axis indicates significance while the x-axis shows the magnitude of expression. Grey dots indicate non-significant genes. Blue dots indicate genes deemed significant yet are lowly expressed. Green dots indicate genes that meet or surpass the log-fold threshold (0.5). Red are genes that are expressed highly and are considered significant. A total of 500 variables were considered. **b)** Dot plot indicating the quantity of nuclei highly expressing genes between NonAD and AD nuclei. Orange = AD samples, Blue = NonAD samples

**Supplementary Figure 3.**
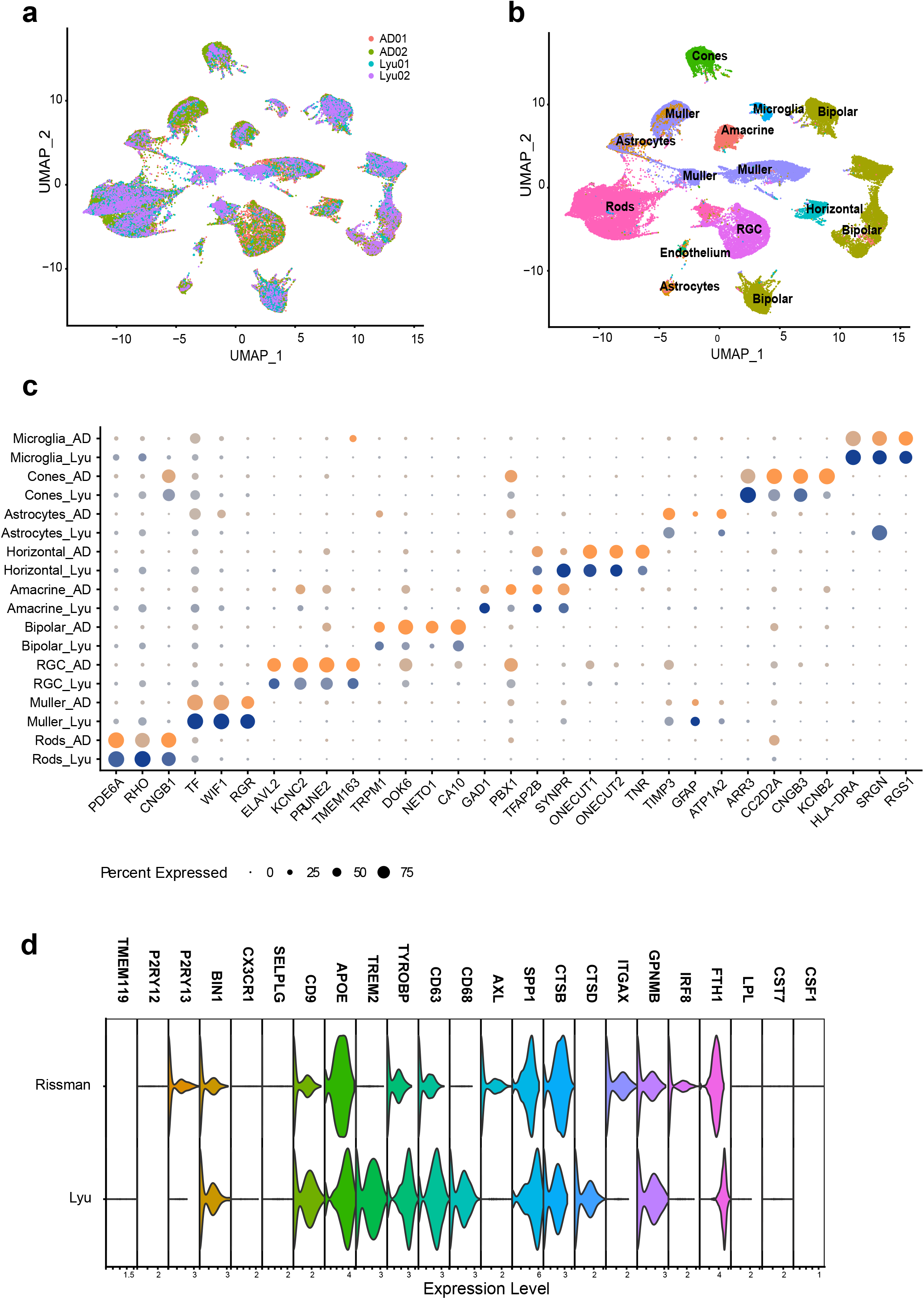
Alignment of single retinal nuclei to single retinal cell dataset indicates potential retinal microglial AD-associated transcriptional signatures. **a)** Clustering of AD retinal nuclei (AD) and non-diseased, elderly retinal cells (Lyu). **b)** Transcriptional expression profiles cluster by retinal cell type. **c)** Dot plot exhibiting single nuclei (orange) and single cell (purple) expression patterns across all cell types. Markers from Figure 1E are used to indicate cell type. **d)** Expression of homeostatic and disease-associated microglial markers between AD and Lyu samples

## Notes

<u>Conflict of Interest:</u> No conflict of interests to report.

### Competing Interest Statement

The authors have declared no competing interest.

## References

1. London, A., Benhar, I. & Schwartz, M. The retina as a window to the brain—from eye research to CNS disorders. Nat. Rev. Neurol. 9, 44–53 (2013).

2. Sinn, R. & Wittbrodt, J. An eye on eye development. Neural Develop. 130, 347–358 (2013).

3. Alber, J. et al. Developing retinal biomarkers for the earliest stages of Alzheimer’s disease: What we know, what we don’t, and how to move forward. Alzheimers Dement. 16, 229–243 (2020).

4. Koronyo, Y. et al. Retinal amyloid pathology and proof-of-concept imaging trial in Alzheimer’s disease. JCI Insight 2, e93621 (2017).

5. Dumitrascu, O. M. et al. Sectoral segmentation of retinal amyloid imaging in subjects with cognitive decline. Alzheimers Dement. Diagn. Assess. Dis. Monit. 12, (2020).

6. Eriksson, U. & Alm, A. Macular thickness decreases with age in normal eyes: a study on the macular thickness map protocol in the Stratus OCT. Br. J. Ophthalmol. 93, 1448 (2009).

7. Chua, J. et al. Age-related changes of individual macular retinal layers among Asians. Sci. Rep. 9, 20352 (2019).

8. Koronyo-Hamaoui, M. et al. Identification of amyloid plaques in retinas from Alzheimer’s patients and noninvasive in vivo optical imaging of retinal plaques in a mouse model. NeuroImage 54, S204–S217 (2011).

9. La Morgia, C. et al. Melanopsin retinal ganglion cell loss in Alzheimer disease. Ann. Neurol. 79, 90–109 (2015).

10. den Haan, J. et al. Amyloid-beta and phosphorylated tau in post-mortem Alzheimer’s disease retinas. Acta Neuropathol. Commun. 6, (2018).

11. Perez, S. E., Lumayag, S., Kovacs, B., Mufson, E. J. & Xu, S. β-Amyloid Deposition and Functional Impairment in the Retina of the APPswe/PS1ΔE9 Transgenic Mouse Model of Alzheimer’s Disease. Investig. Opthalmology Vis. Sci. 50, 793 (2009).

12. Williams, E. A. et al. Absence of Alzheimer Disease Neuropathologic Changes in Eyes of Subjects With Alzheimer Disease. J. Neuropathol. Exp. Neurol. 76, 376–383 (2017).

13. Ho, C.-Y., Troncoso, J. C., Knox, D., Stark, W. & Eberhart, C. G. Beta-Amyloid, Phospho-Tau and Alpha-Synuclein Deposits Similar to Those in the Brain Are Not Identified in the Eyes of Alzheimer’s and Parkinson’s Disease Patients. Brain Pathol. 24, 25–32 (2013).

14. Leger, F. et al. Protein Aggregation in the Aging Retina. J. Neuropathol. Exp. Neurol. 70, 63–68 (2011).

15. Schön, C. et al. Long-Term In Vivo Imaging of Fibrillar Tau in the Retina of P301S Transgenic Mice. PLoS ONE 7, e53547 (2012).

16. Surguchov, A., McMahan, B., Masliah, E. & Surgucheva, I. Synucleins in ocular tissues. J. Neurosci. Res. 65, 68–77 (2001).

17. Grimaldi, A. et al. Neuroinflammatory Processes, A1 Astrocyte Activation and Protein Aggregation in the Retina of Alzheimer’s Disease Patients, Possible Biomarkers for Early Diagnosis. Front. Neurosci. 13, (2019).

18. Ying, P. et al. Single-Cell RNA Sequencing of Retina:New Looks for Gene Marker and Old Diseases. Front. Mol. Biosci. 8, (2021).

19. Lukowski, S. W. et al. A single-cell transcriptome atlas of the adult human retina. EMBO J. 38, e100811 (2019).

20. Liang, Q. et al. Single-nuclei RNA-seq on human retinal tissue provides improved transcriptome profiling. Nat. Commun. 10, 5743 (2019).

21. Olah, M. et al. Single cell RNA sequencing of human microglia uncovers a subset associated with Alzheimer’s disease. Nat. Commun. 11, 6129 (2020).

22. Velmeshev, D. et al. Single-cell genomics identifies cell type-specific molecular changes in autism. Science 364, 685–689 (2019).

23. Mathys, H. et al. Single-cell transcriptomic analysis of Alzheimer’s disease. Nature 570, 332–337 (2019).

24. Gerrits, E., Heng, Y., Boddeke, E. W. G. M. & Eggen, B. J. L. Transcriptional profiling of microglia; current state of the art and future perspectives. Glia 68, 740–755 (2020).

25. Morsey, B. et al. Cryopreservation of microglia enables single-cell RNA sequencing with minimal effects on disease-related gene expression patterns. iScience 24, 102357 (2021).

26. Thrupp, N. et al. Single-Nucleus RNA-Seq Is Not Suitable for Detection of Microglial Activation Genes in Humans. Cell Rep. 32, 108189 (2020).

27. Lyu, Y. et al. Implication of specific retinal cell-type involvement and gene expression changes in AMD progression using integrative analysis of single-cell and bulk RNA-seq profiling. Sci. Rep. 11, 15612 (2021).

28. Yi, W. et al. A single-cell transcriptome atlas of the aging human and macaque retina. Natl. Sci. Rev. 8, nwaa179 (2021).

29. Menon, M. et al. Single-cell transcriptomic atlas of the human retina identifies cell types associated with age-related macular degeneration. Nat. Commun. 10, 4902 (2019).

30. Caglayan, E., Liu, Y. & Konopka, G. Ambient RNA analysis reveals misinterpreted and masked cell types in brain single-nuclei datasets. bioRxiv 2022.03.09.483658 (2022) doi:10.1101/2022.03.09.483658.

31. Leak, R. K. Heat shock proteins in neurodegenerative disorders and aging. J. Cell Commun. Signal. 8, 293–310 (2014).

32. Pottier, C. et al. TYROBP genetic variants in early-onset Alzheimer’s disease. Neurobiol. Aging 48, 222.e9–222.e15 (2016).

33. Keren-Shaul, H. et al. A Unique Microglia Type Associated with Restricting Development of Alzheimer’s Disease. Cell 169, 1276–1290.e17 (2017).

34. Audrain, M. et al. Reactive or transgenic increase in microglial TYROBP reveals a TREM2-independent TYROBP–APOE link in wild-type and Alzheimer’s-related mice. Alzheimers Dement. 17, 149–163 (2021).

35. Lake, B. B. et al. A comparative strategy for single-nucleus and single-cell transcriptomes confirms accuracy in predicted cell-type expression from nuclear RNA. Sci. Rep. 7, 6031 (2017).

36. Wu, H., Kirita, Y., Donnelly, E. L. & Humphreys, B. D. Advantages of Single-Nucleus over Single-Cell RNA Sequencing of Adult Kidney: Rare Cell Types and Novel Cell States Revealed in Fibrosis. J. Am. Soc. Nephrol. 30, 23 (2019).

37. Swamy, V. S., Fufa, T. D., Hufnagel, R. B. & McGaughey, D. M. Building the mega single-cell transcriptome ocular meta-atlas. GigaScience 10, giab061 (2021).

38. Cimino, P. J. et al. Ablation of the microglial protein DOCK2 reduces amyloid burden in a mouse model of Alzheimer’s disease. Exp. Mol. Pathol. 94, 366–371 (2013).

39. Hansen, D. V., Hanson, J. E. & Sheng, M. Microglia in Alzheimer’s disease. J. Cell Biol. 217, 459–472 (2017).

40. Reiman, E. M. et al. GAB2 Alleles Modify Alzheimer’s Risk in APOE ε4 Carriers. Neuron 54, 713–720 (2007).

41. Albrecht, N. E. et al. Rapid and Integrative Discovery of Retina Regulatory Molecules. Cell Rep. 24, 2506–2519 (2018).

