## Supplementary material for "Cell-associated Transcriptional Alterations in the Retinal of Alzheimer’s Disease": Description of Supplemental Files

**Description of Supplementary Figures and Tables**

**Supplementary Figure 1.** Transcriptional characterization of retinal regions between AD and NonAD single retinal nuclei.  **A)** Volcano plot of differentially expressed genes between fovea and periphery. The y axis indicates significance while the x axis shows the magnitude of expression. Grey dots indicate non-significant genes. Blue dots indicate genes deemed significant yet are lowly expressed. Green dots indicate genes that are meet or surpass the log-fold threshold (0.5). Red are genes that are expressed highly and are considered significant. A total of 1759 variables were considered in creating the graph. **B)** Expression of regional-specific genes between AD and NonAD samples. Blue = AD samples, Grey = NonAD samples.

**Supplementary Figure 2.**  Slight dysregulation of gene expression within AD samples. **A)** Volcano plot of differentially expressed genes between NonAD and AD samples. The y axis indicates significance while the x axis shows the magnitude of expression. Grey dots indicate non-significant genes. Blue dots indicate genes deemed significant yet are lowly expressed. Green dots indicate genes that are meet or surpass the log-fold threshold (0.5). Red are genes that are expressed highly and are considered significant. A total of 500 variables were considered. **B)** Dotplot indicating quantity of nuclei highly expressing genes between NonAD and AD nuclei. Orange = AD samples, Blue = NonAD samples

**Supplementary Figure 3.** Alignment of single retinal nuclei to single retinal cell dataset indicate potential retinal microglial AD-associated transcriptional signatures. **A)** Clustering of AD retinal nuclei (AD) and non-diseased, elderly retinal cells (Lyu). **B)** Transcriptional expression profiles cluster by retinal cell type. **C)** Dotplot exhibiting single nuclei (orange) and single cell (purple) expression patterns across all cell types. Markers from Figure 1E are presented to represent each cell type. **D)** Expression of homeostatic and disease-associated microglial markers between AD and Lyu samples.

**Supplementary Table 1.** Retinal marker genes used for cluster assignment, differentially expressed genes in all 31 clusters and gene enrichment analysis within cell types. Data relevant to Figure 1.

**Supplementary Table 2.** Differential gene expression analysis between fovea and peripheral regions. Data relevant to Supplementary Figure 1

**Supplementary Table 3.** Differential expression between AD and NonAD samples. Data relevant to Supplementary Figure 2.

**Supplementary Table 4.** Expression of AD-specific genes between AD, NonAD and Lyu samples. Data relevant to Supplementary Figure 3.
